# Design principles of small intestinal crypt maintenance

**DOI:** 10.1101/2024.07.19.604296

**Authors:** Isidora Banjac, Martti Maimets, Ingrid Tsang, Marius Dioli, Stine Lind Hansen, Kata Krizic, Raul Bardini Bressan, Cecilia Lövkvist, Kim B. Jensen

## Abstract

The intestinal epithelium has a remarkably high turnover in homeostasis. It remains unresolved how turnover is orchestrated at the cellular level and how the behaviour of stem and progenitor cells ensures tissue maintenance. To address this, we combined quantitative fate mapping in three complementary mouse models with mathematical modelling and single-cell RNA sequencing. Our integrated approach generated a spatially and temporally defined model of crypt maintenance that is based on two cycling populations: crypt-based columnar (CBC) and transit amplifying (TA) cells. Validation experiments substantiated the predictions from the model revealing TA cells as the major contributor to the absorptive lineage, while balanced CBC cell fate choices controlled the numbers of cells in the secretory lineage. By unravelling these mechanisms, we gain insights into the process of tissue turnover and provide direct evidence to support the notion of CBC cells as the major driver of the intestinal epithelium replenishment.

**Highlights:**

- Spatially and temporally resolved cellular model of small intestinal crypt maintenance.
- Stem cells exhibit predisposition towards secretory fate.
- Secretory lineage has a limited contribution to the high turnover of intestinal epithelium.
- Cellular differentiation within the crypt occurs rapidly within 72 hours.

## Introduction

The intestinal epithelium has a fast cellular turnover rate and requires continuous production of large numbers of cells^1^. To maintain a healthy and operating tissue, this process must be tightly regulated both in terms of overall numbers as well as proportions of each functional cell type. Along the length of the small intestine, the epithelium is stereotypically organised into crypts containing proliferating cells and villi decorated with postmitotic differentiated cells including absorptive enterocytes, as well as secretory enteroendocrine, goblet, and tuft cells. At the crypt bottom, crypt-based-columnar (CBC) cells, important players in tissue replenishment, are intercalated between secretory Paneth cells ^2,3^. Their progeny, including additional proliferative cells, moves upwards to the top of the crypt and onto the villi while differentiating into the absorptive or secretory lineage ^4,5^.

CBC cells have been proposed to go through an intermediate progenitor state before committing to either of the lineages^1^. Cells expressing early markers of secretory fate, such as Atoh1 and Dll1, appear in positions just above the crypt bottom^5–7^, although it is unclear if these represent progenitors or already maturing cells. In parallel, absorptive cells arise from TAs located in the upper part of the crypt^8^, but specific markers to investigate their behaviour have not been identified yet. Although there are reports of divergent mitotic activities in the crypt^8^, it is not entirely clear how frequent these progenitors divide, and how they contribute to the tissue turnover. During steady state, the final output of turnover is biased towards absorptive lineage, constituting approximately 90% of the epithelium^9^. However, the underlying cause of this difference remains unclear. It could be attributed to the varying proliferative capabilities of the progenitors, biased cell fate choices of CBC cells, or a combination of both. Although, fate mapping studies have provided insights into the turnover of CBC cells^10,11^, a comprehensive model describing general cell behaviour in the crypt is currently lacking.

Our understanding of the design principles in biological systems is significantly enhanced using mathematical modelling approaches. Several studies shed light onto intricate behaviour of stem cells^10–14^, including an attempt to reconstruct spatial patterns of signalling cues for proliferation and differentiation in adult crypts^15^. However, a significant gap remains in understanding the principles governing the short-term maintenance of the intestinal crypt and how cell fate choices balance proliferative activity to maintain a stable turnover in homeostasis.

Here, we employed a multidisciplinary approach to unravel the cellular mechanisms that mediate the homeostatic maintenance of the murine small intestinal epithelium. We propose a model that comprises two spatially separated proliferative populations with distinct cell cycle times. To validate our model, we conducted clonal analyses using two different mouse models, performed EdU/BrdU double labelling experiments and single-cell RNA sequencing (scRNA-seq). Our findings demonstrate that during homeostasis, CBC cells and absorptive progenitors actively contribute to cellular turnover, while secretory precursors do not as their emergence coincides with a postmitotic state.

## Results

### *Lrig1* a marker of the lower crypt epithelium

The crypt epithelium comprises a functionally heterogeneous population of cells with unique roles and divergent proliferative capabilities. To characterize the cellular heterogeneity, we performed scRNA-seq profiling of 2,125 individual crypt cells isolated from the distal small intestine (Ileum). Based on known cell-type markers^16^, cells were clustered into seven transcriptionally distinct populations corresponding to enterocytes, enteroendocrine (EECs), goblet, Paneth, CBC, TA and tuft cells (Figure 1A-C and S1). As expected, the TA and CBC cell clusters displayed the highest expression of cell cycle signature genes, while we also observed smaller fractions of cells with comparably high score in both the enterocyte and goblet cell clusters (Figure 1D).

**Figure 1:**
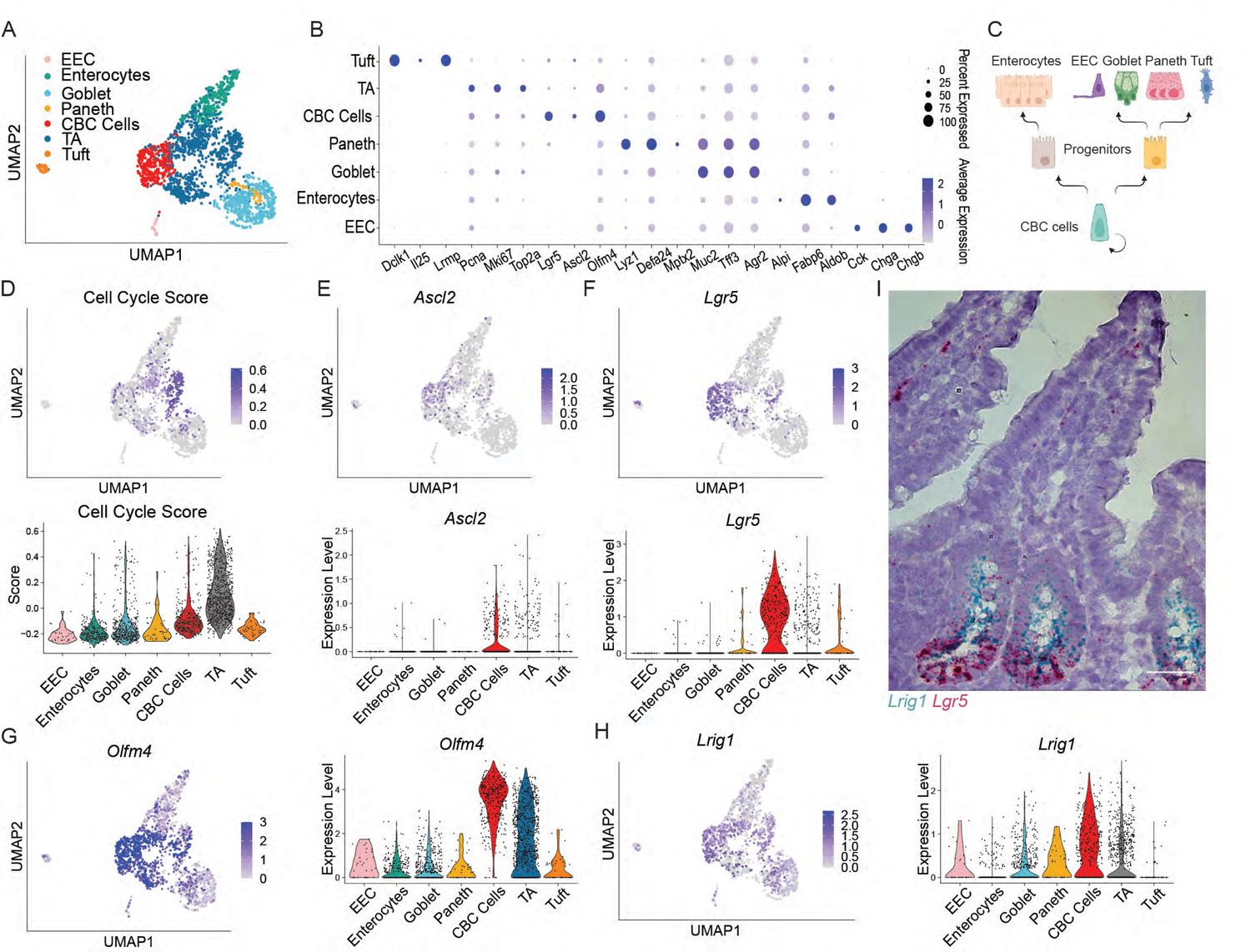
Lrig1 encompasses different populations in the crypt. A) UMAP representing clusters of cell types present in mouse distal small intestinal crypts. B) Dot plot representing expression of known marker genes^16^ used for annotation of the clusters. C) Schematics of the cellular hierarchy in the small intestinal crypt. D) UMAP representing the cell cycle gene signature score; Violin plot representing the cell cycle gene signature score in the clusters. E-H) Normalised *Ascl2, Lgr5, Olfm4, Lrig1* expression in each cluster on UMAP and violin plot. I) In situ hybridisation for *Lrig1* (blue) and *Lgr5* (red) in the distal small intestine. Scale bar, 40um.

To functionally test the contribution of different cell populations to crypt maintenance, including TA and CBC cells as well as secretory cells, we sought to identify a suitable marker for simultaneously targeting these in the crypt. Markers such as *Ascl2*^17^*, Lgr5*^4^ and *Olfm4*^18^ were detected in the CBC/TA clusters (Figure 1E-G). *Lrig1,* a previously described marker of cells in the lower crypt^19^, was detected in CBC/TA cells as well as in some of the additional crypt clusters from both secretory and absorptive lineages (Figure 1H). Single molecule RNA in situ hybridisation for *Lrig1* and *Lgr5* confirmed a more widespread expression of *Lrig1* beyond the CBC region (Figure 1I). Together this data shows that *Lrig1* is a suitable marker for unbiased monitoring of multiple proliferative cell populations in the crypt.

### The existence of two distinct patterns of proliferation within the crypt

To analyse the contribution of Lrig1-expressing cells to crypt maintenance, we performed clonal fate mapping analysis using the *Lrig1-EGFP-IRES-CreERT2*;*Rosa26-mTmG* (Lrig1;mTmG) mouse model. Individual Lrig1-expressing cells were labelled with low-dose tamoxifen and tissue collected for up to 4 days post-labelling (Figure 2A).

**Figure 2:**
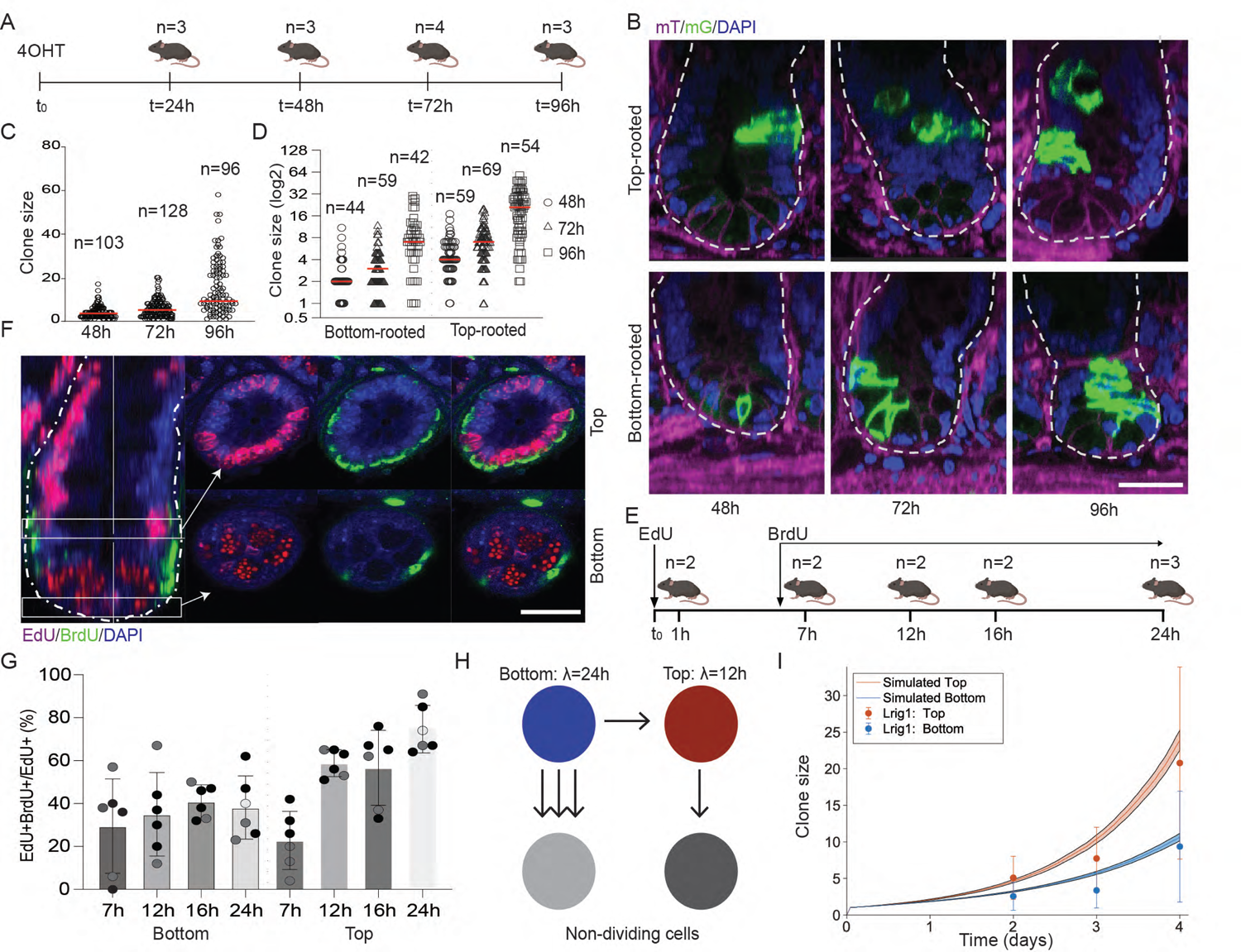
There are two distinct types of behaviour in the crypt. A) Schematics of the fate mapping experiment using Lrig1;mTmG model. Following labelling tissue was collected at 24h intervals until 96h post-injection. B) Confocal microscopy images showing xz/yz rendering of representative top and bottom rooted Lrig1 clones at 48h, 72h and 96h. Membranous signal from clone cells is green (mG) and non-clone cells magenta (mT). Nuclei stained with DAPI (blue). Scale bar, 20um. C-D) Lrig1 clone sizes over time and segregated based on position (top or bottom-rooted). Each point represents one clone; the number of measurements is indicated and the red line represents the mean. E) Schematics of the EdU/BrdU double labelling experiment. F) Confocal microscopy images showing xy/xz rendering and xy view of a representative crypt, EdU (magenta), BrdU (green), DAPI (blue). Scalebar, 20um. G) Percentage of EdU+BrdU+ cells out of EdU+ for bottom and top compartment. Each point represents one crypt (coloured by mice), and error bars represent mean and standard deviation. H) Schematics of the mathematical model. I) Mean clone sizes of top-(red) and bottom-rooted (blue) clones. The model predictions are represented by lines and the shaded areas represent the 95% confidence intervals. Experimental data are means ± s.d. Model parameters are listed in supplementary table 2.

Consistent with *Lrig1* mRNA expression pattern (Figure 1H-I), the first labelled cells were detected at the bottom and higher up in the crypt (Figure 2B) with average clone sizes increasing over time (Figure 2C). To probe whether there are differences in clonal behaviour at the bottom and top of the crypt, labelled clones were classified as either bottom- or top-rooted (Figure S2A). Bottom-rooted clones contained at least one cell located at the bottom of the crypt (irrespective of the position of the remaining cells), whereas top-rooted clones were limited to the upper crypt (Figure 2B, S2A). Importantly, frequencies of top- and bottom-rooted clones were close to 50% at all time points (Figure S2B). During the tracing, bottom-rooted clones increased from 2.5 cells at 48h to 9.4 cells at 96h, while top-rooted clones grew from 5.1 to 20.8 cells (Figure 2D). This two-fold difference in clonal size indicates a pronounced increase in the division rates of upper cells compared to CBC cells at the bottom.

To further investigate how cell cycle time differs along the crypt axis, we measured crypt proliferation kinetics using EdU/BrdU double-labelling. EdU was first pulsed, followed with continuous administration of BrdU 6 hours later (Figure 2E). Tissue was analysed 7, 12, 16 and 24 hours after the initial EdU pulse, and the single- and double-labelled cells quantified in the two compartments at each time point (Figure 2E, F). In the top compartment, 23±6% (mean ± S.D.) of EdU+ cells were double labelled at 7h (Figure 2G). This number increased significantly, plateauing at around 60% between 12h and 16h and rose again to 75±11% by 24h. This strongly suggested that the average cell cycle time was between 12 and 16h. In contrast, in the bottom compartment, EdU labelling was observed in 30±22% of the cells at 7h and plateaued at around 40% from 12h onwards. This demonstrated that the cell cycle kinetics is on average slower in the lower compared to the upper compartment (Figure 2G), aligning with an observation that the Lgr5-expressing CBC cells cycle around once per 24h^10^.

### Mathematical modelling of intestinal crypt turnover

As cells within the crypt transition from the bottom to the top, some bottom cells naturally assume the position of top cells. Since the division times differ between these compartments, the contribution of a cell to clone growth changes accordingly. Fate mapping experiments provide a snapshot of the clone at a given moment, without reference to prior positions of its cells and their behaviour. Therefore, it becomes challenging to draw definitive conclusions related to contributions from bottom versus top cells and compartment-specific cell cycle times. Based on previous successful use of mathematical modelling to investigate biological paradigms^10–14^, we next applied this framework to the generated fate mapping data to develop a model for crypt cell behaviour.

We first tested if the clonal dynamics could be explained by two populations (top and bottom) representing independent sources of turnover. Here crypt maintenance is driven by two independent populations of continuously dividing cells with longer and shorter cell cycle. A cell cycle length of λ=24h was assigned to bottom cells, while a range of shorter cell cycles λ=12-16h was used for the top population based on predictions from the EdU/BrdU double labelling experiment. Clonal labelling and expansion were simulated in this model over 4 days mimicking the Lrig1;mTmG fate mapping. The simulation resulted in two different growth curves for bottom and top clones. When aligned with the results from Lrig1;mTmG lineage tracing, the predicted clone sizes were not matching the experimental data (Figure S2C). A model with two independent cycling populations is therefore insufficient to explain the data.

To test the alternative and account for bottom cells differentiating into top cells, we introduced a probability for this transition. The newly simulated growth curves captured the feature of top clones being almost double the size of the bottom ones (1.6x bigger at 96h), but the predicted clone sizes of both clone types were still out of the range of experimental values (Figure S2D). Thus far, the model did not include any probability to differentiate into post-mitotic cells within the time frame of the simulation. To improve the fit of the model, cells in the top were allowed to differentiate and thereby stop dividing. CBC cells were expected to cycle continuously and only differentiate through proliferative progenitor populations. (Figure S2E). With these modifications to the model, bottom clones in the simulation grew larger than the top clones, which clearly does not fit the experimental data. To test how clones would grow if cells at the bottom had the possibility to differentiate to postmitotic cells, we introduced a non-dividing bottom cell. This provided bottom cells with two choices, as they could either become a top cycling cell with a probability (p) or stop dividing (1-p). By making the choice for a bottom cell to stop dividing a preferred choice (1-p=0.7), while restricting the cell cycle of top cells to 12h (Figure 2H, Figure S2F-G), the model accurately predicted the clone sizes of both bottom and top-rooted clones (Figure 2I).

In summary, the Lrig1;mTmG fate mapping data can be explained by 1) different cell cycle times in the top and bottom compartment (Figure 2G) and 2) direct differentiation of both bottom and top cells into post-mitotic differentiated cells. Importantly, the mathematical model suggests a bias for bottom cells to transition into postmitotic rather than cycling top cells (Figure 2H).

### Lgr5+ cells at the crypt bottom function as a homogeneous population

CBC cells expressing Lgr5 have been shown to be equipotent stem cells that self-renew with each division giving rise to stem cells that compete for space in the niche^10,11^. To assess if the behaviour of the CBC cells can be explained by the proposed model, we performed clonal fate mapping using *Lgr5-EGFP-IRES-CreERT2*;*Rosa26-mTmG* (Lgr5;mTmG) mouse model (Figure 3A-D). Tissue was collected for 3D imaging at 24h intervals up to 4 days post labelling. At 24h, most labelled cells were positioned at the crypt bottom (83%), with the remainder observed right above, at positions 4 and 5 (Figure 3B, S3A). Over time, clones grew rapidly and expanded towards the top compartment, increasing the frequency of top-rooted clones (Figure 3B, C). However, when clones were segregated into top and bottom-rooted groups, we did not observe any significant size difference (Bottom: 1.7 cells at 24h to 12.8 at 96h vs Top 1.3 cells to 15.2) (Figure 3D).

**Figure 3:**
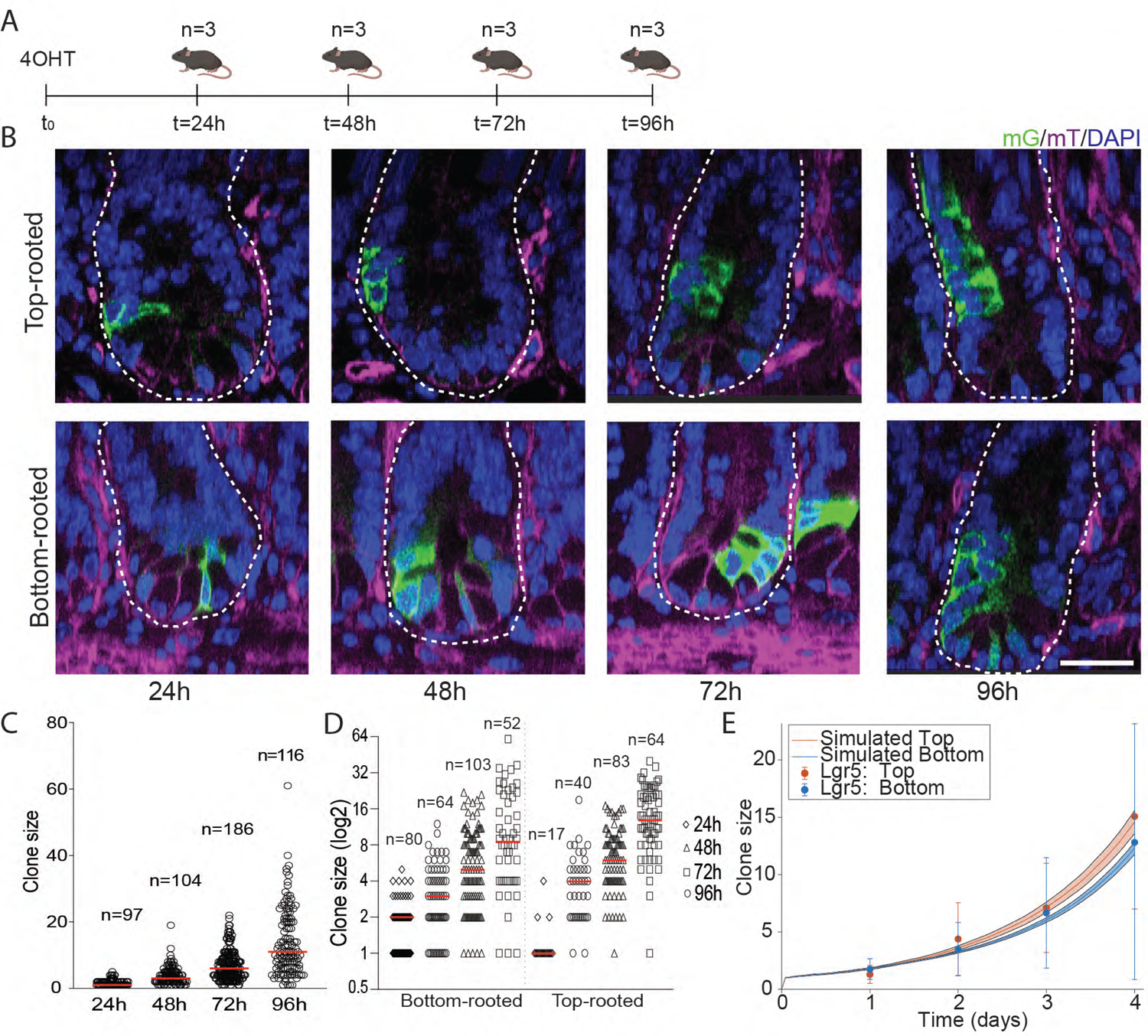
Lgr5+ cells at the bottom behave in a homogeneous manner. A) Schematics of the fate mapping experiment using Lgr5;mTmG model. Tissue was subsequently collected at 24h intervals until 96h post-injection. B) Images showing a xz/yz rendering of 3D imaged representative top- and bottom-rooted Lgr5 clones at 48h, 72h and 96h green (mG) and magenta (mT). Nuclei stained with DAPI (blue). Scale bar, 20um. C-D) Lgr5 clone sizes over time and segregated based on their position (top- or bottom-rooted). Each point represents one clone; the number of measurements is indicated and the red line represents the mean. E) Mean Lgr5 clone sizes of top (red) and bottom (blue) rooted clones. The model predictions are represented by lines and the shaded areas represent the 95% confidence intervals. Experimental data are means ± s.d. Model parameters are listed in supplementary table 2.

To challenge the predictive capacity of our mathematical model (Figure 2H), we simulated labelling of CBC cells using the parameters inferred from the Lrig1;mTmG experiment (Figure 2H). Evidently, the model has strong predictive powers as both bottom and top-rooted clone sizes are predicted with a high degree of accuracy (Figure 3E). This strongly supports a model where CBC cells undergo symmetrical cell divisions while their progeny can either continue dividing as CBCs, transition into the upper crypt compartment or differentiate directly into postmitotic cells. Moreover, the model predicts a preference for this fate choice over moving into the fast-cycling top cell compartment.

### Non-dividing cells are present in the clones at 48h and 72h

Intrigued by the possibility of Lgr5+ CBC cells directly committing to differentiation, we analysed the cellular composition of Lgr5;mTmG progeny 48h and 72h after labelling via scRNA-seq (Figure 4A). Non-recombined epithelial cells were isolated as an internal control representing the heterogeneity of the intestinal epithelium (Figure S4A-C). Overall, we profiled 1,371 labelled cells that were clustered and annotated as before (Figure 1A-C; 4B). The cells were further separated into progeny from the 48h (171 cells) and 72h samples (1,199 cells). The analysis showed that progeny from the 48h samples consisted mostly of CBC cell/TA compartment (40%/40%), together with enterocytes, goblet, Paneth and EECs, but no tuft cells (Figure 4B, S4D). Strikingly, the 72h sample contained all the cell types detected in the control epithelium (Figure 4B), with differentiated cells represented in higher proportions than in the 48h, but not reaching those detected in the control samples (Figure 4B, S4D). The increase of differentiated cell types in the 72h clones demonstrated that the differentiation process from an Lgr5+ CBC cell to any differentiated lineage occurred within as little as 72h.

**Figure 4:**
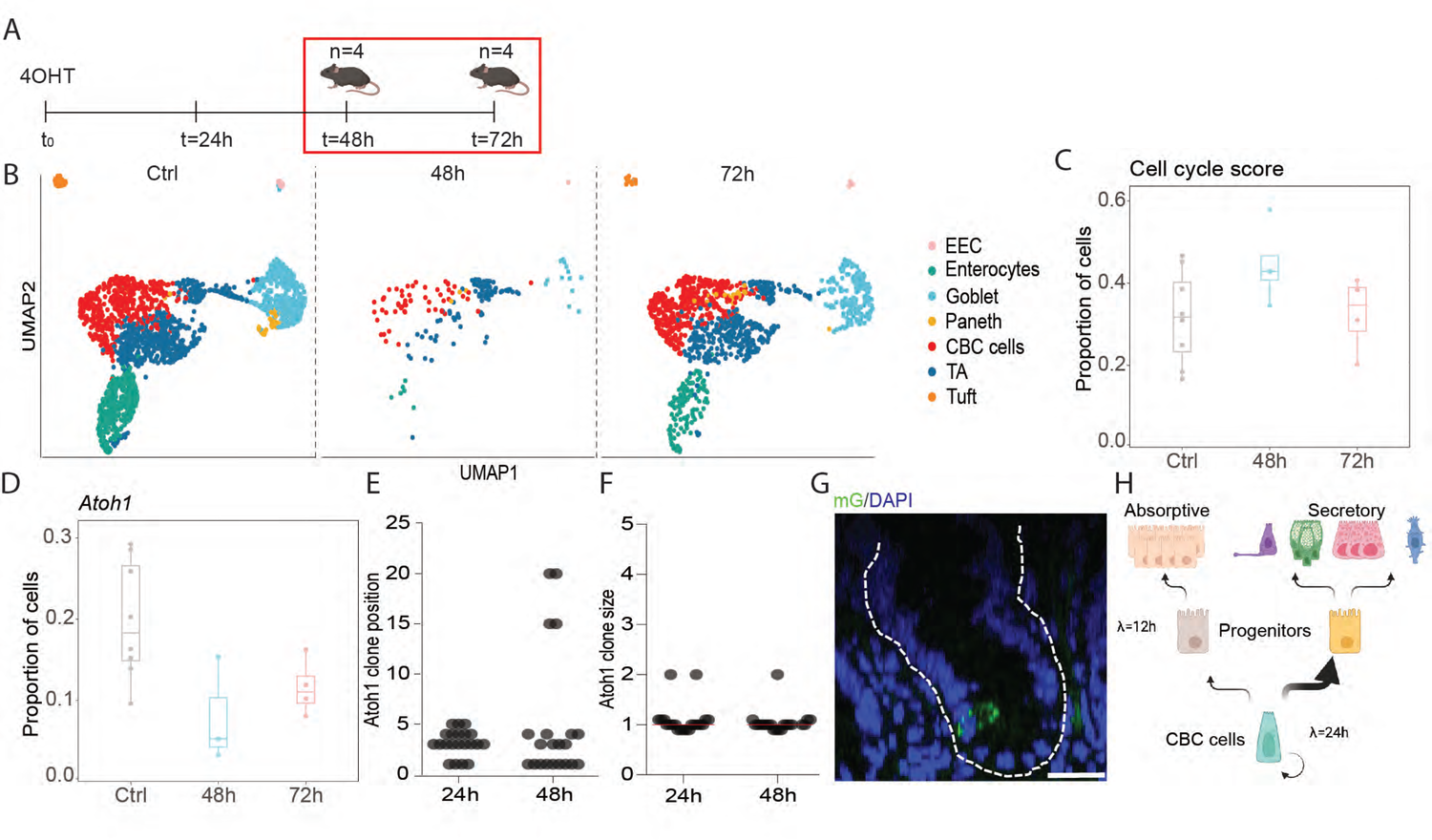
Secretory cells do not contribute significantly to the crypt maintenance. A) Schematics of the single cell RNA sequencing experiment using Lgr5;mTmG model. Following labelling with mG tissue was subsequently collected at 48h and 72h post and cells prepared for single cell RNA sequencing. B) UMAP representing clusters of cell types present in mouse distal small intestinal crypts in control tissue and clones at 48h and 72h. C-D) Proportion of proliferative cells based on cell cycle genes signature score and *Atoh1* expressing cells within each sample. Boxplots show the medians, and the dots represent the samples. E-F) Clone sizes and position of Atoh1 clones 24h and 48h post labelling. Each point represents one clone. G) Side view of 3D imaged representative Atoh1 clone at 48h. Green (mGFP) blue (DAPI). Scale bar, 20um. H) Proposed model of homeostatic maintenance of distal small intestinal epithelium. 1-cell cycle time.

As our model proposed that Lgr5+ CBC cells had the capacity to differentiate directly to at least one cell type, we wondered if there was evidence for this in the scRNA-seq data^4^. Interestingly, transcriptional analysis revealed that on average 25% of CBC cells in the crypt epithelium expressed *Lyz1*, a marker of Paneth cells (Figure S4E)^20^. These observations were further substantiated by detection of Lgr5-EGFP+Lyz1+ cells in the crypt (Figure S4F). This suggested that Paneth cells could differentiate directly from an Lgr5+ CBC cell without passing through a fast-cycling state (Figure S4E, F). Paneth cells are consequently a suitable candidate for a non-cycling cell state predicted by our mathematical model (Figure 2H). This opened the question whether other secretory cells could follow similar patterns, thereby regulating secretory cell numbers by simply adjusting fate choices of CBC cells.

### Limited short-term contribution of Atoh1+ cells to the crypt turnover

Transient upregulation of *Atoh1* is a requirement for differentiation into the secretory lineage^6^. In the scRNA-seq data, 5% of cells at 48h expressed *Atoh1*, increasing to 10% at 72h versus 18% in the control epithelium (Figure 4D). To assess the proliferative capabilities of *Atoh1* expressing cells, we performed fate mapping using the *Atoh1-*CreERT2;Rosa26*-mTmG* mouse model (Atoh1;mTmG) (Figure 4E-G). Labelled cells were initially observed either at the crypt bottom or at position 4-5, in line with previous reports^5^ (Figure 4E). In time, cells either rapidly moved out of the crypt or were retained at the crypt bottom (Figure 4E). Importantly, the number of labelled cells per crypt remained constant between 24h and 48h (Figure 4F) suggesting that *Atoh1* expressing cells differentiated directly into the secretory lineages without entering a cycling state. These cells therefore do not contribute significantly to the high turnover of the small intestinal epithelium in line with our model.

## Discussion

It is crucial to understand the principles of cellular replenishment of tissues. Here, we have taken an interdisciplinary approach combining quantitative 3D fate mapping, mathematical modelling and single-cell transcriptomics to unravel the complex and dynamic behaviours of cells within the intestinal epithelium. Our model revealed that the epithelium is maintained by two proliferating populations: 1) CBC cells biased towards the secretory lineage and 2) TA daughter cells committed to the absorptive lineage. Whereas the CBC cells undergo cell divisions approximately every 24 hours, the TA progeny displays a faster cell cycle estimated at 12 hours.

Importantly, our model predicts that most CBC cells differentiate directly into postmitotic progenitors of the secretory lineage. In contrast, absorptive enterocytes are generated via continued cell division in the TA compartment^3^, thereby accounting for the higher proportion of absorptive cells within the epithelium. The temporally resolved scRNA-seq analysis of Lgr5+ progeny revealed that differentiation to all lineages occurred within 72h, while cells are still located within the crypt. Since the turnover rate for the small intestinal epithelium is estimated at around 3-5 days^2^, the fast pace of differentiation is necessary to generate enough cells to replace those exfoliated into the lumen from tips of villi. Our model revealed that the observed clonal behaviour could only be achieved through continuous proliferation of CBC cells and TAs, with CBC cells biased towards a non-dividing secretory fate.

Drawing from prior models of cell behaviour in the crypt^10,11^, the simplicity of the proposed model and its reliance on experimentally validated parameters enhances its credibility. Furthermore, predicted clone sizes from our model are in excellent agreement with both Lrig1 (Figure 2I) and Lgr5 (Figure 3E) lineage tracing data. Additionally, the concept of cells exiting cell cycle while they are committing to a postmitotic secretory fate, has been proposed before and aligns well with the results of the fate mapping data from Atoh1 expressing cells^22,23^. Retrograde movement of cells into the CBC compartment during homeostasis has been proposed^21^. As it is challenging to align our clonal fate mapping data with the model encompassing substantial retrograde movement, we conclude this is a rare event that does not contribute significantly to the homeostatic maintenance.

An additional advantage of our approach is the definition of different cell types based on their proliferative behaviour, allowing us to analyse their emergence and proportions within the clones. This proves particularly beneficial since specific markers for the TA population are lacking, making their independent analysis challenging. By taking a reductionistic approach and subtracting the behaviours of cell types with known markers, such as Lgr5+ (CBC cells) and Atoh1+ (secretory progenitors), we can deduce the behaviour of the TAs. This strategy provides valuable insights into the dynamics of the intestinal crypt and potentially other proliferative tissues where distinct markers for all the cell types are not known. Interestingly, Lrig1 has previously been proposed to be a marker of quiescent stem cells in the intestinal epithelium^24^. However, in line with other observations^19^, both our single cell expression and fate mapping clearly demonstrate that Lrig1 is expressed by proliferating cells in the crypt and that these cells are actively contributing to tissue homeostasis.

As more definitive information on cell populations in the crypt emerges, our model can be updated and refined, ensuring its adaptability and relevance to current knowledge. This valuable tool advances intestinal modelling approaches and deepens our understanding of the complex behaviour of intestinal crypts.

### Limitation of study

Defining cell types primarily based on their proliferative behaviour could potentially oversimplify the complexity of the cellular landscape. While recognising the importance of proliferation in defining cell function, we acknowledge that it is not the sole determinant of cellular identity. By categorising cells into discrete states based on proliferation, we have not considered other crucial characteristics that contribute to cell behaviour and function, including signalling pathways, gene expression profiles, and microenvironmental cues. Emerging views of cell differentiation propose a more continuous and dynamic representation of cellular states^1^, challenging the rigid categorization used in this study^1^. Incorporating additional factors and exploring the continuum of cell states could provide a more comprehensive understanding of the system and its underlying complexities.

## Acknowledgments

We thank members of the Jensen laboratory for technical assistance and comments on the manuscript, as well as the imaging, flow cytometry and genomics/bioinformatics facilities at reNEW and BRIC, and animal caretakers for expert assistance as well as Oliver Knights Møller from the Single-Cell Omics platform at the Novo Nordisk Foundation Center for Basic Metabolic Research (CBMR) for the technical expertise and support. We kindly thank Doug J. Winton for sharing with us the Atoh1-creERT2 mouse line and Mariaceleste Aragona for critical discussions. Work in the Jensen lab is supported by the Independent Research Fund Denmark (0134-00111B to K.B.J.), Novo Nordisk Foundation (NNF20OC0064376 to K.B.J.), Leo Pharma Foundation (LF-OC-19-000169). I.B. is the recipient of a fellowship from the Novo Nordisk Foundation as part of the Copenhagen Bioscience Ph.D. Program (NNF18CC0033666). RBB is supported by a Marie Skłodowska-Curie postdoctoral fellowship (895802/H2020-MSCA-IF-2019). The Novo Nordisk Foundation Center for Stem Cell Medicine is supported by a Novo Nordisk Foundation grant (NNF21CC0073729). Figure elements were adapted from BioRender.

## Author Contributions

Conceptualisation: I.B., C.L., K.B.J.; Methodology: I.B., M.M., S.L.H., C.L., K.B.J.; Validation: I.B., C.L., K.B.J.; Formal Analysis: C.L., I.T., M.D., I.B.; Investigation: I.B., M.M., S.L.H., K.K., R.B.B.; Data Curation: I.B., C.L., K.B.J.; Writing-Original Draft: I.B., C.L.; Writing-Review & Editing: M.M., K.B.J.; Visualisation: I.B., C.L., M.M., K.B.J.; Supervision: M.M.,C.L., K.B.J.,; Project Administration: I.B., C.L., K.B.J.; Funding Acquisition: K.B.J.

## STAR methods

### Resource availability

#### Lead contact

Further information and requests for resources and reagents should be directed to and will be fulfilled by the Lead Contact, Kim Bak Jensen.

#### Materials availability

This study did not generate new unique reagents.

### Experimental model and subject details

#### Mice

Unless otherwise specified C57BL/6J mice (purchased from Taconic, Denmark) were used for the experiments. Transgenic murine lines used in the experiments have been previously described: *Lgr5-EGFP-IRES-CreERT2*^4^*, Lrig1-EGFP-IRES-CreERT2*^25^, *Atoh1-CreERT2*^26^ (kind gift from D. Winton) and *Rosa26-mTmG*^27^. The mice were housed under controlled conditions in specific-pathogen-free (SPF) facilities. Food and water were provided ad libitum. Both male and female mice (except C57BL/6J that were only females) aged 8-11 weeks were used in the experiments, distributed into randomised cohorts for each experimental timepoint. All animal procedures had been reviewed and approved by The Danish animal inspectorate (Permit number 2023-15-0201-01378).

## Method details

### Lineage tracing

For *Cre* induction animals received single intraperitoneal (*i.p.)* injection of 4-OH-tamoxifen (H6278, Sigma-Aldrich) (dissolved in ethanol and mixed with corn oil). Dose titration was performed for each transgenic line to achieve single cell labelling per crypt: 80ug/20g body weight for Lrig1;mTmG, 100ug/20g body weight for Lgr5;mTmG and 20ug/20g mouse for Atoh1;mTmG. Lgr5; mTmG animals were labelled with 1.5mg/20g body weight for isolation of clones for scRNA-seq.

### EdU/BrdU proliferation experiments

For the double pulse experiments, mice were first injected *i.p.* with EdU (5-ethynyl-2’-deoxyuridine) (A10044, Invitrogen™/ Thermo Fisher Scientific) as a DPBS solution 200ug/20g body weight. Secondly, BrdU (5-bromo-2’-deoxyuridine) (11594167, Invitrogen™) was administered 6h later *i.p.* as a DPBS solution 2mg/20g body weight as well as in the drinking water 0.8mg/mL with 1% glucose. Subsequently, mice were sacrificed 1h, 7h, 12h, 16h and 24h after the first injection for sample collection.

### Intestinal whole-mount and immunostaining

Mice were sacrificed by cervical dislocation and 5 cm of distal small intestine (measured from caecum) was collected, flushed with ice cold PBS, opened longitudinally, washed again, and fixed overnight in 4% PFA on 4°C.

Tissue for wholemount imaging was then cut into pieces of 1cmx1cm and cleared while shaking on 37°C in CUBIC-1A (5% N,N,N′,N′-Tetrakis(2-hydroxypropyl) ethylenediamine (122262, Sigma-Aldrich), 10% Urea (U5378, Sigma-Aldrich), 10% Triton X-100 (X100, Sigma-Aldrich), 25mM NaCl in PBS) with DAPI (4’,6-diamidino-2-phenylindole) (D9542, Sigma-Aldrich) for 5-7 days, refreshing the solution every second day. 30min before imaging tissue was mounted on the glass slides in RapiClear 1.52 (RC152002, Sunjin Lab) and imaged within a week.

Immunostaining for lysozyme was performed after the clearing step, by first washing the tissue in PBS and then incubating in 1%BSA 0.5%Triton PBS with primary Ig Rabbit polyclonal anti-human Lysozyme EC 3.2.1.17 (AB_2341230, Agilent Dako) 1:500 for 48h, shaking on +4°C. After washing with PBS, it was incubated in 1%BSA 0.5%Triton PBS with secondary Donkey anti-rabbit 647 Ig (AB_2536183, ThermoFisher Scientific) for additional 48h, shaking on +4°C.

EdU/BrdU staining was performed on the wholemounts right after fixation. Wholemounts were first incubated 30min in preheated 37°C HCl, then EdU signal was developed according to the manufacturer’s instructions (Click-iT® EdU Alexa Fluor® 647 Imaging Kit, C10340, ThermoFisher Scientific). Finally, BrdU staining was performed by first incubating the tissue in the blocking buffer (1%BSA (A9576, Sigma-Aldrich), 5% Goat Serum (G9023, Sigma-Aldrich), 0.8% Triton X-100 (X100, Sigma-Aldrich) in PBS) overnight, then incubated with the primary anti-BrdU antibody (AB_626519, Santa Cruz) 1:50 in blocking buffer for 2 days on room temperature and then incubated for 2h with the secondary Goat anti-mouse IgG2a 555 (AB_2535776, ThermoFisher Scientific) 1:200 antibody on room temperature. Tissue was then cleared and mounted as described above.

For in situ hybridisation tissue was processed according to manufacturer’s protocols. Briefly, following fixation the intestinal pieces were incubated in 10%, 20% and 30% sucrose (1h each) and embedded in OCT for cryosectioning. 10um sections were cut on the Leica Cryostat and co-ISH was performed using RNAscope 2.5 Duplex Reagent Kit (322430, ACD Biotechne) by the instructions of the manufacturer with probes Mm-Lgr5 (312171-C2, ACD Biotechne) and Mm-Lrig1 (310521,ACD Biotechne).

### Imaging and Image analysis

3D images of the crypts were acquired using laser-scanning confocal microscopy (Leica TSC SP8 or Leica Stellaris Fluorescence Confocal microscope) with a Z-step of 1-3um. Sections with ISH were acquired with Leica DM5500 Widefield. Images were imported in Imaris (SCR_007370, Bitplane, Oxford, US) and quantified manually.

### Isolation of single cells for scRNA-seq

Mouse distal small intestine (2x 3cm) was collected, flushed with ice cold PBS and opened, while villi were scraped with a coverglass. After an additional wash, tissue was incubated in HBSS Ca+Mg+ Free (14175095, ThermoFischer Scientific) containing 5mM EDTA (15575-020, Invitrogen) 10mM HEPES (15630-056, Gibco/Life Technologies) for 25 min at 37°C, vortexing every 5 min. Crypts were shaken out, washed with PBS and then incubated in TrypLE (12605036, ThermoFisher Scientific) supplemented with 10uM y27632 (Y0503, Sigma-Aldrich), 1mM NAC (A9165, Sigma-Aldrich), 50ug/ml DNaseI (10104159001, Roche) for 21min at 37°C and pipetted 50x every 5min to dissociate into single cells. Cells were then pelleted in aDMEM/F12 (12634-028, Gibco/Life Technologies) containing 5% FBS, washed in 0.1% BSA/PBS, strained through 100um cell strainer, washed again in 0.1% BSA/PBS and filtered through blue-capped FACS tubes. Subsequently, cells were stained in 0.1% BSA/PBS, containing following antibodies: EpCAM-APC (AB_1659714, eBioscience, 1:500) CD31 PeCy7 (AB_10612003, BD biosciences, 1:200), CD45 PeCy7 (AB_394489, BD biosciences, 1:200), DAPI 1ug/mL, Hashing Ig (1:100) and transferred to low retention tubes, incubated 15min in the dark on room temperature, washed with 0.1%BSA and resuspended in ultra clean 0.1% BSA/PBS for flow cytometry. Samples were sorted with BD FACS Aria III sorter, 70u nozzle, 1.0 ND filter into an Eppendorf tube with 2uL ultra clean 0.1% BSA/PBS.

Hashing was performed so that each sample contained cells from one mouse and was labelled by a distinct hashing Ig before the sort. This means we had 8 hashed samples from 4 mice that were traced 48h (# 1-4) and 4 mice traced 72h (# 5-8). We sorted GFP^high^ clones and GFP-cells from the epithelium as control into 2 eppendorfs so that each contained cells from 48h and 72h clones, as well as the control epithelium, and each contained all 8 hashes. Thus, we had 2 samples for library preparation: Sample 1 contained control epithelium from mice number 1,2,5,6, 48h clones from mice 3,4 and 72h clones from mice 7,8; Sample 2 contained control epithelium from mice number 3,4,7,8, 48h clones from 1,2 and 72h clones from 5,6.

### Preparation of libraries and single cell RNA sequencing

The libraries were prepared using the Chromium Next GEM Single Cell 3’ Reagent Kits v3.1 according to the 10X Genomics protocols v3.1. Maximum 10,000 cells were loaded into 36uL of ultra clean 0.1%BSA/PBS for the formation of Gel Bead-in-Emulsions and followed by reverse transcriptase. cDNA was amplified with 12 polymerase chain reaction (PCR) cycles, while sample-indexing PCR was performed with 14 cycles. The libraries were sequenced on an Illumina NextSeq 2000 platform with a High Output 150 cycles kit.

### Processing of raw scRNA-seq reads

The 10x Genomics Cell Ranger 6.1.2 software was used to process the scRNA-seq output and to generate the count matrices used for the downstream analysis. The refdata-gex-mm10-2020-A reference was downloaded from the 10x Genomics website (https://support.10xgenomics.com). The FASTQ files were generated by the Cell Ranger *mkfastq* function and further aligned and filtered using the Cell Ranger *count* function. This process was performed on both libraries and generated filtered feature-barcode matrices. All downstream analysis was performed using Seurat (v.4.3.0)^28^.

### Cell hashing and assignment of samples

The outputted filtered feature-barcode matrices contained both a gene expression matrix (RNA) and an antibody capture matrix (HTO). Cells overlapping between these two matrices was further used. To assign a sample to each cell the HTO counts was first normalised using *NormalizeData* (with normalisation method = ‘CLR’ and margin = 2). For each HTO, the normalised HTO data is then modelled as a mixture of two normal distributions (a negative cluster from background noise and a positive cluster from true HTO signal) using the *mclust* R package ^29^. Cells with higher number of normalised HTO read count than that of the negative HTO cluster are assigned with the corresponding HTO label. By modelling each HTO independently, the cell hashing antibody affinity variation is considered. Cells are finally demultiplexed into sample identity, where cells without HTO assignment are labelled as negative, with only one HTO assignment as singlets and with more than one HTO assignment as doublets. We further proceed with cells labelled as singlets and as most doublets are removed during this process, we did not perform any additional doublet filtering.

### Quality control and filtering of cells

Low quality cells were filtered out by excluding cells that have unique feature counts over 6000 or less than 200. Additionally, cell with less counts than 500 were also filtered out. The percentage of mitochondrial gene counts per cell was calculated using *PercentageFeatureSet* and cells with more than 50% mitochondrial counts were filtered out. Moreover, the log10 transform of the number of genes per UMI for each cell was calculated, and cells with a value below 0.8 were filtered out. To further remove technical effects, the genes *Malat1*, *Gm42418* and mitochondrial genes were filtered out.

### Merging libraries and analysis of CTRL and Lgr5 clones

The datasets were further analysed in two ways. First the CTRL samples (GFP-cells) were pulled out from both libraries and analysed together. Secondly, all samples were analysed together to analyse the Lgr5 clones (GFP^high^ cells) and compare to the CTRL samples. The two types of analysis are described in the following sections.

### Control samples

The cells from the control samples of each sequence library, were separated by sample (HTO) and then each sample where were normalised using the function *NormalizeData*, this log transformed for each cell by total counts over all genes. To integrate the datasets, the function *FindVariableFeatures* was first used on each sample. Then, the samples were integrated used *FindIntegrationAnchors* including anchor features from *SelectIntegrationFeatures*. The samples were finally integrated using *IntegrateData* and have thereby batch corrected over the HTOs.

The S phase score and G2M phase score was calculated using *CellCycleScoring* on the cell cycle genes provided by Seurat. The scores were then regressed out using *ScaleData* along with the mitochondrial counts. The scaling was done on all genes in the dataset. The overall *Cell Cycle Score*, used to assess the proliferative cells, was calculated as a combined score based on all cell cycle genes provided by Seurat using the function *AddModuleScore*.

### Lrg5 clones

To analyse the Lgr5 clones (GFP^high^ cells) together with the CTRL (GFP-cells), the two libraries were merged using the function merge. The combined dataset was then normalised and log transformed for each cell by total counts over all genes (using *NormalizeData*). The *FindVariableFeatures* was run on the full dataset. The S phase score and G2M phase score was calculated using *CellCycleScoring*, similar to the control samples. Then the difference between the S phases score and the G2M phase was regressed out using *ScaleData*, and the scaling was done on all genes in the dataset.

### Clustering, dimensionality reduction and cluster annotations

Dimensionality reduction was performed using principal-component analysis (*RunPCA*) followed by Uniform Manifold Approximation and Projection (*RunUMAP*). To further cluster the cells, the functions *FindNeighbors* and *FindClusters* was used. For clustering and UMAP analyses 13 (Lgr5 clones + CTRL) and 14 (CTRL) principal components were used. The expression of known markers^16^(Figure 1B and S1) guided the cell type annotations of a resolution of 7 clusters. The same procedure was performed on both the CTRL data and the combined CTRL and Lgr5 clone dataset.

### Cell proportion plots

When proportions of cells expressing a selected gene or the cell cycle score, are calculated, a cell is considered to express a gene if the scaled expression is above 0. For each timepoint (48h and 72h) and for the CTRL, the proportion of expressing cells are calculated within each sample.

### Simulations of the mathematical model

To simulate the proliferation dynamics and the contribution of different populations to the clones, we used the following modelling scheme. The cells can either be a dividing bottom cell (*B*_d_), a non-dividing bottom cell (*B*_n_), a dividing top cell (*T*_d_) or a non-dividing top cell (*T*_n_) (Figure 2H). The cells in a clone are simulated as independent agents, where each cell have the possibility to either divide or differentiate. Moreover, the two processes are separated from each other^10,11^.

When a cell divides it generates a new cell with the same status, i.e., a *B*_d_ will generate a new *B*_d_ and a *T*_d_ will generate a new *T*_d_. When a bottom cell changes state, it can either become a dividing top cell *B*_d_ → *T*_d_, with a probability *p* or a non-dividing bottom cell *B*_d_ → *B*_n_, with a probability of 1 − *p*. A dividing top cell can become a non-dividing top cell, *T*_d_ → *T*_n_. Additionally, a non-dividing bottom cell move upwards and become a non-dividing top cell, *B*_n_ → *T*_n_.

A simulation of a clone is carried out as following:

The simulation starts off with one cell initialised in one of the four states (*B*_d_, *B*_n_, *T*_d_ and *T*_n_). The initial state is dependent on the type of labelling that is simulated, (see table 2). The simulation is then conducted over t time steps and at each time step the following actions are implemented: For each cell in *B*_d_ and *T*_d_, a random number *r* will be drawn. If *r* < *p*_d_ (see table 1) the cell will divide, and a new cell will be added to the clone with the same status (see above). For each cell in *B*_d_, *B*_n_ and *T*_d_ a random number *r* is drawn and if *r* < *p*_n_^T^, *p*_n_^BT^ or *p*_t_^B^ (depending on the cell’s state) the cell will change state according to table 1.

**Table 1.** Description of division and state change probabilities.

| Parameter | Description |
| --- | --- |
| $p_d$ | $p_d^B$ The probability per time step for a bottom cell to divide. |
| | $p_d^T$ The probability per time step for a top cell to divide. |
| $p_n^T$ | The probability of a $T_d$ to stop dividing per time step (to become $T_n$) |
| $p_n^{BT}$ | The probability of a $B_n$ to move to the top compartment per time step (to become $T_n$) |
| $p_t^B$ | The probability of a $B_d$ to change state per time step (either to $B_n$ or $T_d$) |
| $p$ | The probability of becoming a $T_d$ over a $B_n$ when $B_d$ is changing state. |

**Table 2.** The model parameters used for each figure.

| Parameters | S2C | S2D | S2E | 2I(S2F-G) | 3E |
| --- | --- | --- | --- | --- | --- |
| $p_d^B$ | 0.0289 | 0.0289 | 0.0289 | 0.0289 | 0.0289 |
| $p_d^T$ | 0.0578<br>0.0495<br>0.0433 | 0.0578<br>0.0495<br>0.0433 | 0.0578<br>0.0495<br>0.0433 | 0.0578<br>0.0495 (S2F)<br>0.0433 (S2G) | 0.0578 |
| $p_n^B$ | 0 | 0.0289 | 0.0289 | 0.0289 | 0.0289 |
| $p_n^T$ | 0 | 0 | 0.05 | 0.025 | 0.025 |
| $p_n^{BT}$ | 0 | 0 | 0 | 0.01 | 0.01 |
| $p$ | 0 | 1.0 | 1.0 | 0.3 | 0.3 |
| $w(B_d, B_n, T_d, T_n)$ | (0.7,0,0.3,0) | (0.7,0,0.3,0) | (0.7,0,0.3,0) | (0.68,0.02,<br>0.15,0.15) | (0.9711,<br>0.0289,<br>0,0) |

In total N clones are simulated over t time steps (here N=10000 and t=96) and from those average clone sizes are calculated. From the simulation results, a clone is defined as a bottom rooted clone when at least one cell in the clone is either a *B*_d_ or *B*_n_. All other clones are considered as top rooted clones. The average clone sizes are shown with a 95% confidence interval.

### Initialisations of clones to simulate Lrig1 and Lgr5 clones

The simulation of a clone is initialised in different states depending on the experiment of interest. To simulate a Lgr5 clone the simulation is initialised as a bottom cell (*B*_d_ or *B*_n_). The state is randomly chosen from the two but weighted (see Table 2). To simulate a Lrig1 clone, the clone can both be labelled as a bottom and top cell. However, since Lrig1 is observed to be more expressed at the bottom^19^ there is a higher probability to be labelled at the bottom (see weights in Table 2).

### Quantification and statistical analysis

The number of biological and technical replicates and the number of animals are indicated in figure legends and text. All tested animals were included. Sample size was not predetermined. For all experiments with error bars, the standard deviation (SD) was calculated to indicate the variation within each experiment. Significance was assessed using the tests indicated in the figure legends.

### Quantification of clone size and position

Each clone was assessed separately. The clone size is quantified by counting the number of labelled cells within the crypt. The clone position within the crypt is annotated based on the lowest cell in the clone. Looking at the side view, cells located at the bottom, below the lumen forming level, have the position 1-3. Cells at the lumen forming level are at the position 4. Position of cells located higher up in the crypt was annotated by counting the number of nuclei between position 4 and the observed cell in the side view. In the further analysis clones were categorised into bottom rooted and top rooted group based on the position of the lowest cell. If the clone contains a cell that is localised in the positions 1-3 the clone will be classified as bottom rooted one, otherwise it will be a top rooted clone.

### Quantification of EdU/BrdU double labelling experiment

Six crypts (from 2 mice) were analysed for each timepoint in the experiment (except 24h where we analysed crypts from 3 mice). Bottom and the top compartment were quantified separately in each crypt. Cells below the lumen forming level (cells in position 1-3) were analysed as bottom, while cells in the positions 5-8 as top compartment. We excluded cells in position +4. The total number of EdU+, BrdU+, EdU+BrdU+ and unlabelled cells was counted for each compartment per crypt.

---

**Figure S1.**
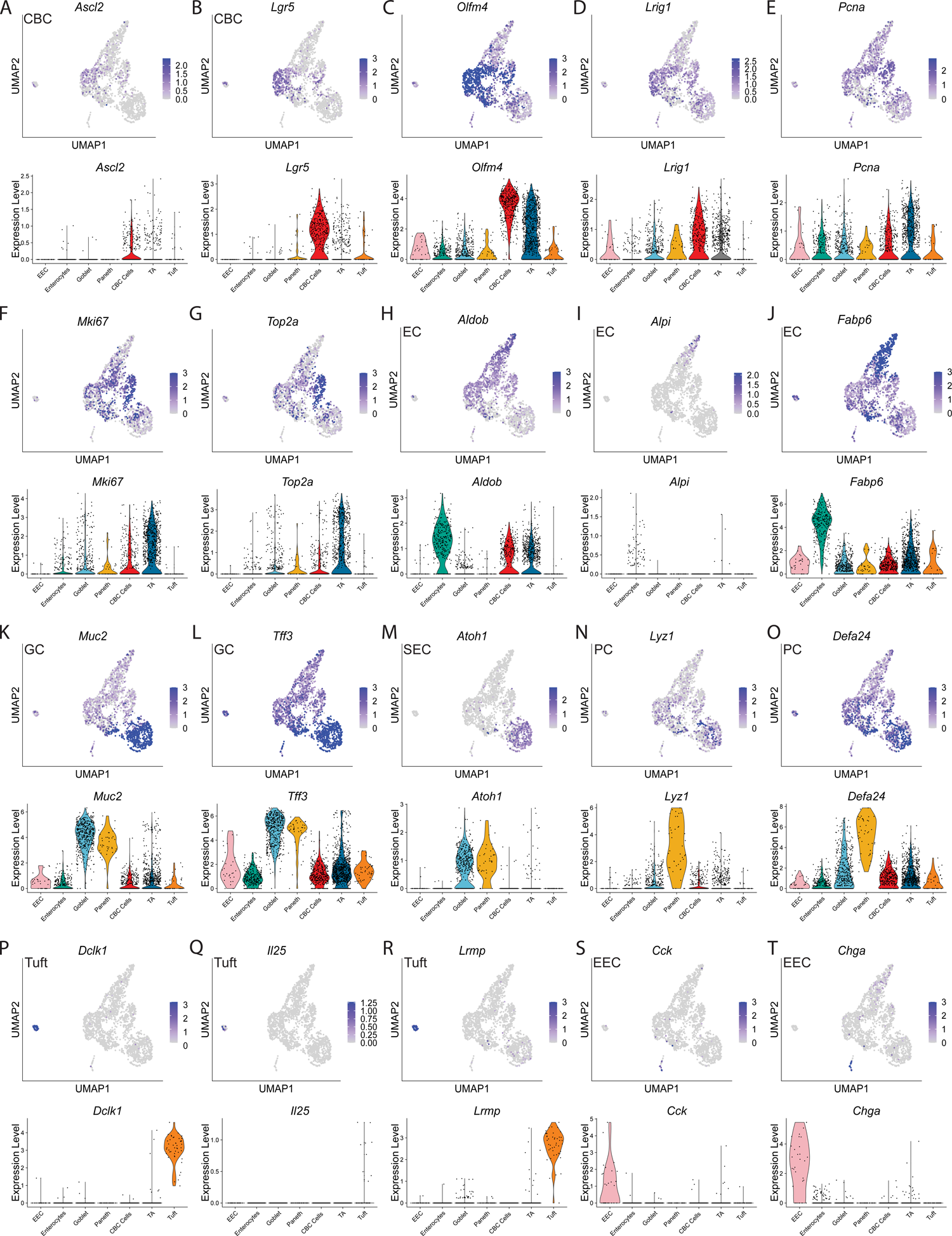
Identifying intestinal epithelial cell-types in scRNA-seq data according to the known marker expression (A) Normalized *Ascl2* expression on UMAP plot; Violin plot of normalized *Ascl2* expression in each cluster. (B) Normalized *Lgr5* expression on UMAP plot; Violin plot of normalized *Lgr5* expression in each cluster. (C) Normalized *Olfm4* expression on UMAP plot; Violin plot of normalized *Olfm4* expression in each cluster. (D) Normalized *Lrig1* expression on UMAP plot; Violin plot of normalized *Lrig1* expression in each cluster. (E) Normalized *Pcna* expression on UMAP plot; Violin plot of normalized *Pcna* expression in each cluster. (F) Normalized *Mki67* expression on UMAP plot; Violin plot of normalized *Mki67* expression in each cluster. (G) Normalized *Top2a* expression on UMAP plot; Violin plot of normalized *Top2a* expression in each cluster. (H) Normalized *Aldob* expression on UMAP plot; Violin plot of normalized *Aldob* expression in each cluster. (I) Normalized *Alpi* expression on UMAP plot; Violin plot of normalized *Alpi* expression in each cluster. (J) Normalized *Fabp6* expression on UMAP plot; Violin plot of normalized *Fabp6* expression in each cluster. (K) Normalized *Muc2* expression on UMAP plot; Violin plot of normalized *Muc2* expression in each cluster. (L) Normalized *Tff3* expression on UMAP plot; Violin plot of normalized *Tff3* expression in each cluster. (M) Normalized *Atoh1* expression on UMAP plot; Violin plot of normalized *Atoh1* expression in each cluster. (N) Normalized *Lyz1* expression on UMAP plot; Violin plot of normalized *Lyz1* expression in each cluster. (O) Normalized *Defa24* expression on UMAP plot; Violin plot of normalized *Defa24* expression in each cluster. (P) Normalized *Dclk1* expression on UMAP plot; Violin plot of normalized *Dclk1* expression in each cluster. (Q) Normalized *Il25* expression on UMAP plot; Violin plot of normalized *Il25* expression in each cluster. (R) Normalized *Lrmp* expression on UMAP plot; Violin plot of normalized *Lrmp* expression in each cluster. (S) Normalized *Cck* expression on UMAP plot; Violin plot of normalized *Cck* expression in each cluster. (T) Normalized *Chga* expression on UMAP plot; Violin plot of normalized *Chga* expression in each cluster. (A-T) CBC-crypt based columnar; EC-enterocyte; GC-goblet cell; SEC-secretory cell; PC-Paneth cell; EEC-enteroendocrine cell.

**Figure S2.**
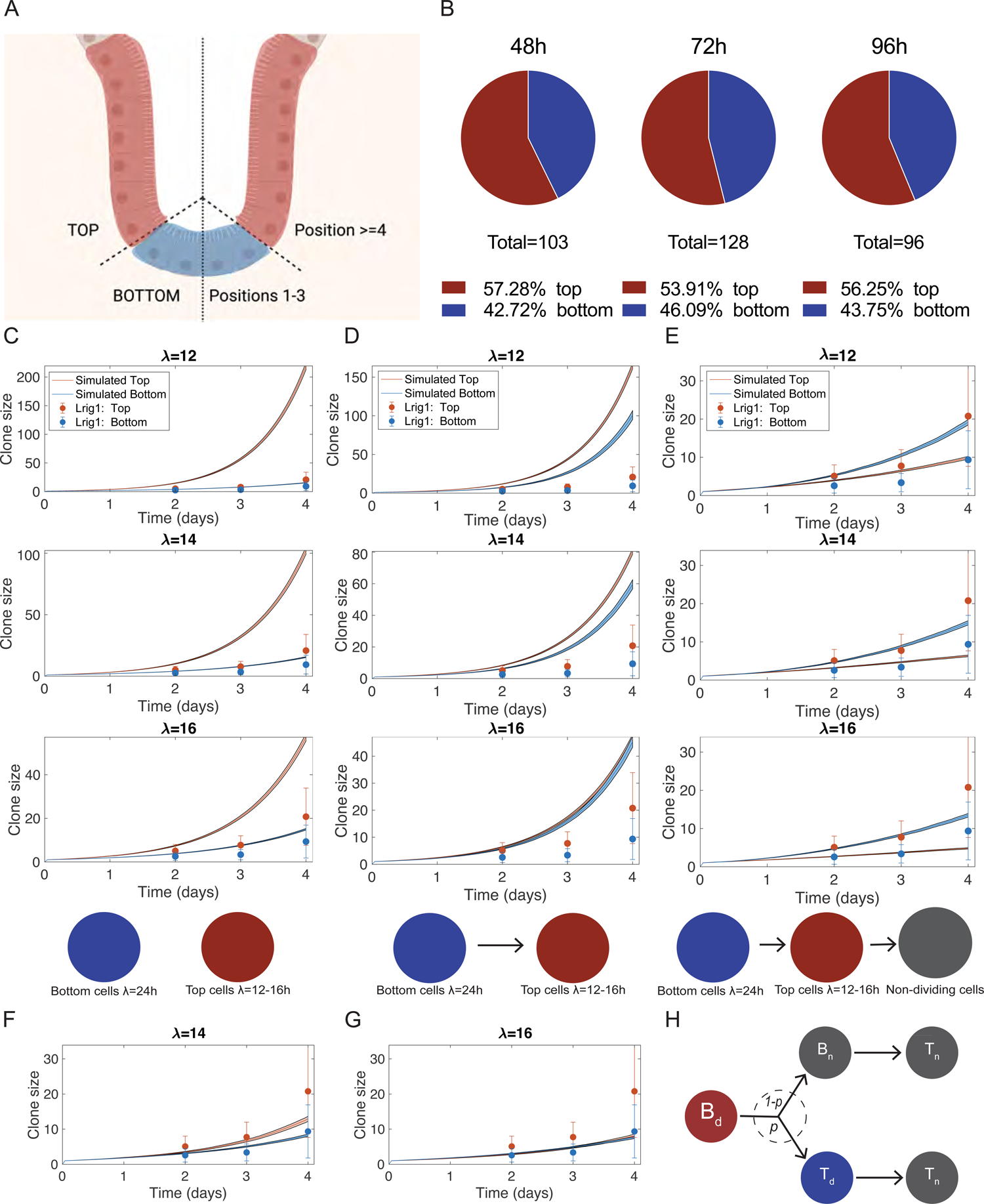
Modelling of the Lrig1; mTmG data. (A) Schematics of the cellular positions in the small intestinal crypt. (B) Distribution of top-rooted and bottom-rooted clones in the Lrig1; mTmG fate mapping experiment. (C) Schematics of the mathematical model and predictions from the model. The model includes two independent populations: bottom cells with a cell cycle (λ) of 24h and a faster cycling top cell with a cell cycle time in the range 12-16h. (D) Schematics of the mathematical model and predictions from the model. The model includes bottom cells with a cell cycle (λ) of 24h that can become a faster cycling top cell with a cell cycle time in the range 12-16h. (E) Schematics of the mathematical model and predictions from the model. The model includes bottom cells with a cell cycle (λ) of 24h that can become a faster cycling top cell with a cell cycle time in the range 12-16h. Top cells have a probability to stop cycling. (F) Predictions of the mathematical model. The model includes bottom cells with a cell cycle (λ) of 24h that can become a faster cycling top cell with a cell cycle time of 14h. Top cells have a probability to stop cycling. (G) Predictions of the mathematical model. The model includes bottom cells with a cell cycle (λ) of 24h that can become a faster cycling top cell with a cell cycle time of 16h. Top cells have a probability to stop cycling. In (C)-(G) mean clone sizes of top (red) and bottom (blue) rooted clones. The model predictions are represented by lines and the shaded areas represent the 95% confidence intervals. Experimental data are means ± s.d. Model parameters are listed in supplementary table 2. (H) Schematics of the mathematical model (Fig 2I and S2F-G) showing the transition from a cycling bottom cell to a cycling top cell with a probability *p* or a postmitotic bottom cell with a probability *1-p*. The cycling top cells can further stop dividing and the postmitotic bottom cells can move and become postmitotic top cells.

**Figure S3.**
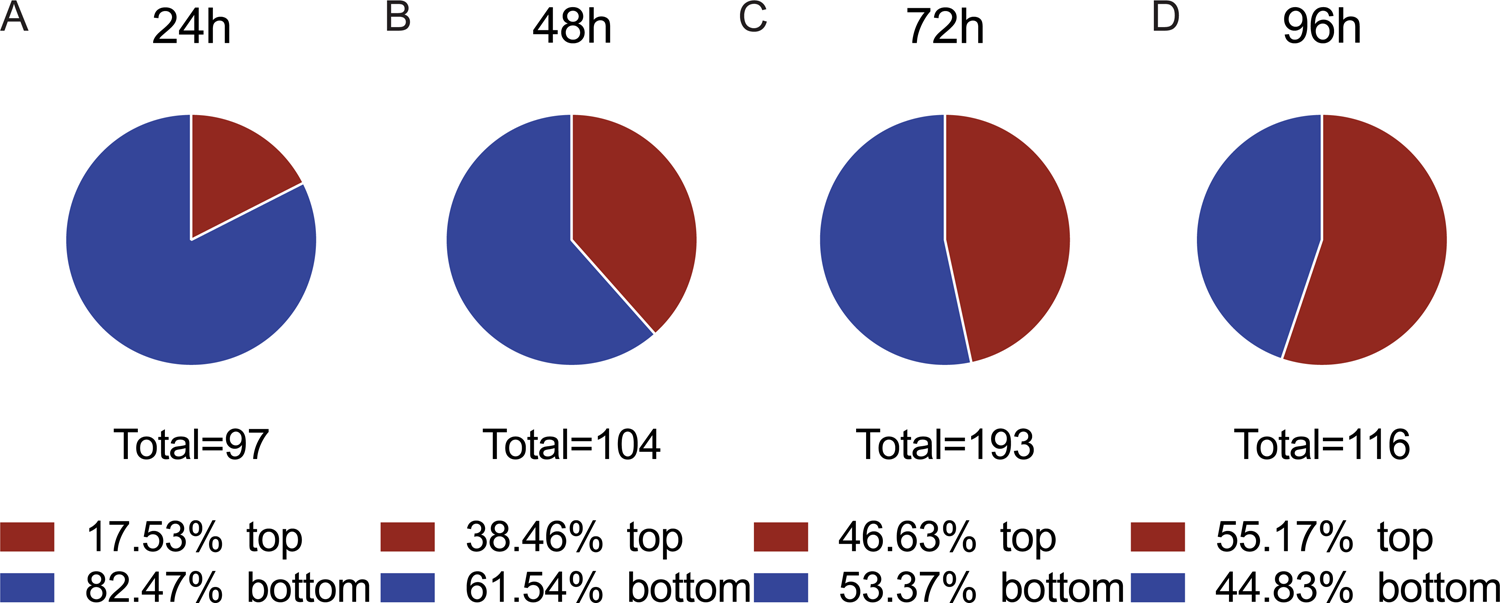
Distribution of top-rooted and bottom-rooted clones. (A) Distribution of top-rooted and bottom-rooted clones in the Lgr5; mTmG fate mapping experiment at 24h. (B) Distribution of top-rooted and bottom-rooted clones in the Lgr5; mTmG fate mapping experiment at 48h. (C) Distribution of top-rooted and bottom-rooted clones in the Lgr5; mTmG fate mapping experiment at 72h. (D) Distribution of top-rooted and bottom-rooted clones in the Lgr5; mTmG fate mapping experiment at 96h.

**Figure S4.**
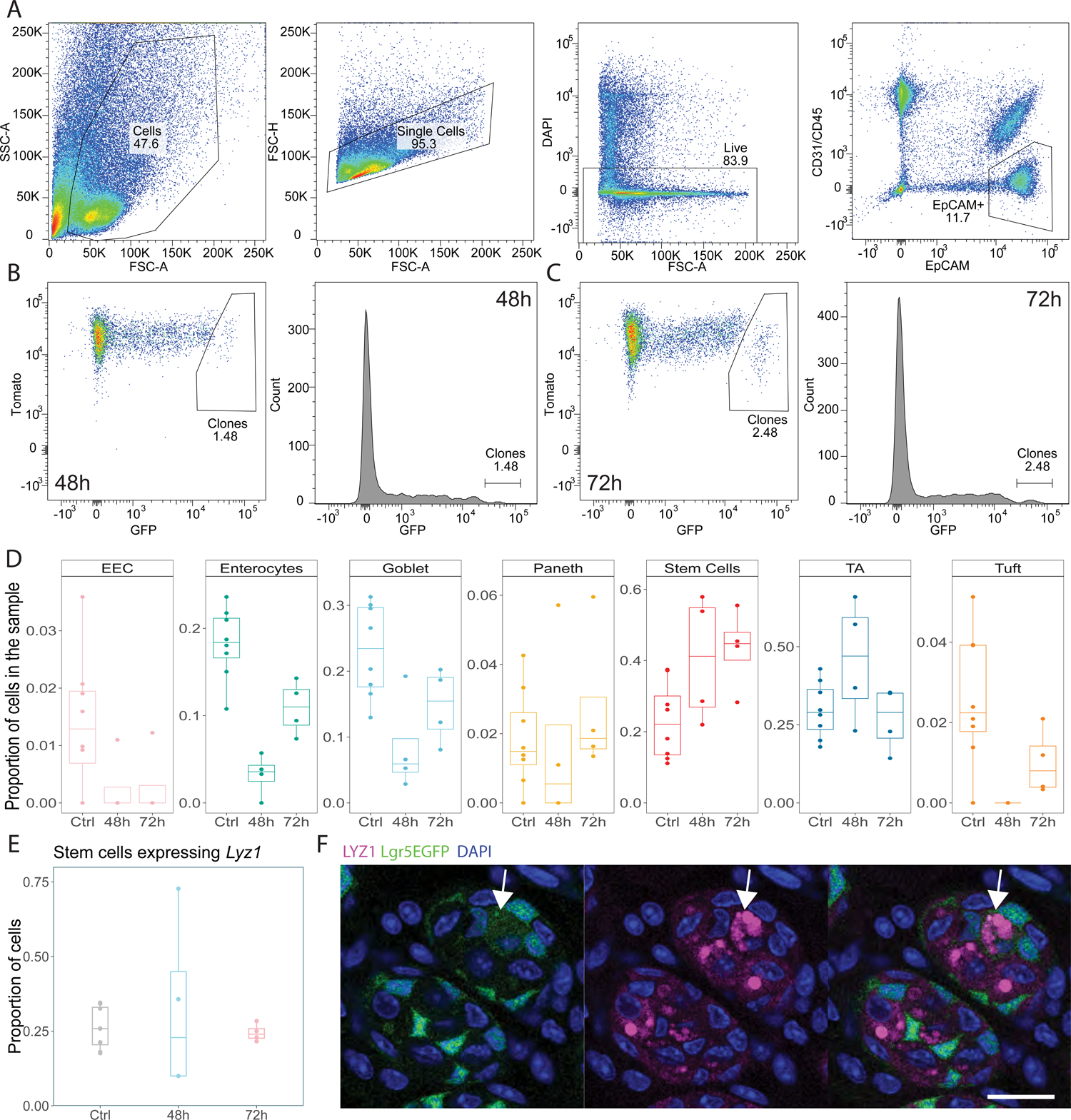
Lgr5+ cells could directly differentiate towards Paneth cells (A) Gating strategy for sorting epithelial cells for scRNAseq (B) Gating strategy for separating clone cells from the rest of the epithelium in 48h sample (C) Gating strategy for separating clone cells from the rest of the epithelium in 72h sample (D) Proportion of annotated clusters within the 48h, 72h and control sample. (E) Proportion of *Lyz1+* expressing cells among stem cells. (F) Confocal microscopy images showing co-localization of Lyz1 antibody (magenta) and Lgr5EGFP(green). Nuclei stained with DAPI (blue). Scale bar, 20um.

